# Regulation of fatty acid oxidation involved in high altitude hypoxia induced cardiomyocyte ferroptosis via the SIRT1-PPARα-GPX4 signaling pathway

**DOI:** 10.1101/2025.01.20.633884

**Authors:** Xiaona Song, Jie Chen, Hongbao Xu, Ling Zhang, Zirou Wang, Lingling Pu, Zhaoli Chen, Xinxing Wang, Weili Liu

**Affiliations:** Tianjin Institute of Environmental and Operational Medicine, Tianjin 300050, China; Department of Taopu Township Community Health Center, Putuo District, Shanghai 200331, China; NO.983 Hospital of PLA Joint Logistics Support Force, Tianjin 300142, China

**Author notes:** Correspondence: Weili Liu, No.1 Dali Road, Heping District, Tianjin 300050, China;.

**Keywords:** Hypobaric hypoxia, Cardiac injury, Fatty acid β-oxidation, PPARα, Ferroptosis, GPX4

## Abstract

The heart experiences damage and undergoes changes in energy metabolism under condition of hypoxia. Ferroptosis is a novel form of cell death characterized by the accumulation of reactive oxygen species (ROS) and excessive lipid oxidation. However, the relationship between cardiac energy metabolism and ferroptosis at high altitudes remains unexplored. This study investigates the effects of high-altitude hypoxia on cardiac function and explores underlying mechanisms. The results demonstrate that exposure to high-altitude hypoxia induces structural and functional damage to the heart, leading to myocardial cell injury and oxidative stress, which are accompanied by ferroptosis. Inhibition of ferroptosis with Fer-1 significantly enhances hypoxia-induced expression of GPX4 and SLC7A11, reduces ROS levels, restores the GSH/GSSG ratio, and improves viability of hypoxia-damaged myocardial cells. Further investigation revealed that hypoxia induces mitochondrial damage and disrupts fatty acid β-oxidation (FAO), accompanied by downregulation of PPARα. Activation of PPARα with WY14643 enhances FAO, suppresses ferroptosis, and boosts cell viability and ATP levels under hypoxic conditions. Overexpression of SIRT1 upregulates PPARα, enhances FAO, and mitigates ferroptosis, these effects are reversed by the PPARα inhibitor GW6471. Resveratrol, a natural SIRT1 activator, improves cardiac function and mitochondrial structure, enhances FAO, and reduces ferroptosis in high-altitude hypoxia-induced cardiac injury through the SIRT1-PPARα-GPX4 pathway. These findings identify the SIRT1-PPARα-GPX4 pathway as a potential therapeutic target for high-altitude-induced cardiac injury.

## 1. Introduction

The defining characteristic of high-altitude environments is persistent hypobaric hypoxia (HH). Despite the atmospheric oxygen content remaining at 21% at all altitudes, ascending leads to reduced barometric pressure and subsequently lowered inspired PO_2_, posing challenges in oxygen delivery to tissues^[1]^. Studies have shown that exposure to HH can induce cardiac hypertrophy, which may potentially lead to cardiac systolic dysfunction and ventricular dilation in severe cases^[2, 3]^. Ferroptosis is an iron-dependent form of cell death triggered by lipid peroxide accumulation and membrane integrity disruption^[4]^. Unique to ferroptosis, compared to apoptosis, necroptosis, and other forms of non-apoptotic cell death, is its central involvement of iron-dependent accumulation of lipid reactive oxygen species (ROS)^[5]^. The lipid hydroperoxidase glutathione peroxidase 4 (GPX4) converts lipid hydroperoxides to lipid alcohols, preventing the ferrous formation of toxic lipid ROS dependent on ferrous iron (Fe^2+^)^[6]^. Recent studies have closely linked ferroptosis with various cardiovascular diseases^[7–9]^, while proteomic analyses of high-altitude populations suggest alterations in transferrin, which is crucial for deiron signaling^[10]^. However, the relationship between high-altitude hypoxia-induced cardiac injury and ferroptosis remains unexplored.

Ferroptosis arises from iron-dependent phospholipid peroxidation, regulated by biological lipid metabolism and associated cellular processes^[11]^. Essentially, ferroptosis represents a consequence of cellular metabolism where oxygen and iron, integral to metabolic processes, lead to the production of ROS as inevitable byproducts. Failure to effectively neutralize lipid peroxides, especially phospholipid peroxides, leads to a loss of membrane integrity and cell death through ferroptosis^[12]^. In recent years, GPX4 has been reported to be associated with PPARα^[13, 14]^. PPARα suppresses ferroptosis by promoting GPX4 expression and inhibiting the expression of plasma iron carrier TRF^[15]^. PPARα is a nuclear receptor that regulates the expression of multiple genes controlling both fatty acid uptake and oxidation, and it is a key player in the transcriptional regulation of cardiac substrate preference^[16]^. Moreover, PPARα is regulated by SIRT1, which ameliorates Ang II-induced impairment of energy metabolism and mitochondrial damage^[17]^. It has been shown that energy metabolism is strongly associated with heart failure, ischemia/reperfusion, and diabetic cardiomyopathy^[18–20]^. Hypoxia is the driving force that regulates the characteristic metabolic switch from primary FAO in the healthy heart to glucose (GLU) utilization in the failing myocardium^[21]^. This metabolic structure is thought to be advantageous under hypoxia because glucose produces 25-60% more ATP per O_2_ molecule compared to free fatty acids (FFA)^[22]^. Nevertheless, the altered energy metabolism induced by HH can still cause damage to the heart. Researchers investigated changes in energy metabolism in right and left ventricular homogenates from rats living in a hypoxic environment. The main finding was that the oxidative capacity of the left ventricle was diminished by hypoxic adaptation^[23]^. It has been noted that an increasing number of metabolic pathways ultimately converge in the process of ferritin deposition^[24]^. The O_2_-dependent oxidation of carbon fuels in mitochondria produces large amounts of ATP but also generates some ROS^[25]^. Hence, we speculated that ferroptosis occurs due to changes in myocardial energy metabolism under HH. Additionally, hypoxia-induced cardiac injury may be associated with ferroptosis.

This study aims to investigate the role of ferroptosis in mediating cardiac damage induced by high-altitude hypoxia. We comprehensively explore the regulatory mechanisms of ferroptosis influenced by changes in myocardial energy metabolism under HH conditions, ultimately aiming to elucidate how reducing ferroptosis through improved energy metabolism could mitigate cardiac function impairment caused by high-altitude hypoxia.

## 2. Materials and Methods

### 2.1. Chemicals and Reagents

Dulbecco’s Modified Eagle Medium (DMEM), penicillin-streptomycin solution, 0.25% trypsin/EDTA, phosphate-buffered saline (PBS), and fetal bovine serum (FBS) were obtained from Gibco (Grand Island, NY, USA). ATP and MDA assay kits were purchased from Beyotime (Nanjing, China). Free fatty acid (FFA) assay kits were obtained from Jianglai Bioengineering Institute (Shanghai, China). GSH and GSSG assay kits were from Abbkine (California, USA). Acetyl-CoA (A-CoA) assay kits were sourced from Grace Biotechnology (Suzhou, China). Antibodies against PPARα, pyruvate dehydrogenase kinase 4 (PDK4), carnitine palmitoyltransferase 1 (CPT1), peroxisome proliferator-activated receptor gamma coactivator-1 alpha (PGC1α), SIRT1, GPX4, SLC7A11, and β-actin were obtained from Proteintech (Wuhan, China) and Abcam (Cambridge, UK). Secondary goat anti-rabbit IgG antibody was purchased from Bioworld (Nanjing, China). ROS assay kits, Dimethyl sulfoxide (DMSO), BCA Protein Quantification Kit, RIPA lysis buffer, protease inhibitor cocktail, SDS-PAGE Gel Preparation Kit, and other western blotting reagents were obtained from Solarbio (Beijing, China). Cell Counting Kit-8 (CCK-8) was procured from Dojindo (Kumamoto, Japan). Hoechst, tetramethylrhodamine ethyl ester (TMRE), BODIPYTM 581/591, WY14643, GW6471, and Ferrostatin-1 were purchased from MCE (New Jersey, USA). Resveratrol (RSV, purity ≥98%) was obtained from Harveybio (Beijing, China). Unless specified, all other chemicals and reagents were from Sigma-Aldrich (St. Louis, MO, USA).

### 2.2. Animal experiments

Animal studies were carried out in accordance with the guidelines of NIH. Animal experiments are approved by the Animal Care and Use Committee of the Ministry of Environmental and Operational Medical Research (SYXK AMMS-04-2021-03, Beijing, China) and following the Guidelines for the Care and Use of Laboratory Animals. Male Wistar rats (7 weeks old) were procured from Beijing Vital River Laboratory Animal Technology Co., Ltd. A hypobaric chamber simulated a 6000 m altitude environment, exposing rats for 20 hours daily to establish a hypobaric hypoxia cardiac injury model.

The study was designed to prepare, characterize and assess improvements in the protective efficacy of RSV in *in vitro* models of hypobaric hypoxia-induced hypertrophy. Rats were divided into three groups: Control, hypobaric hypoxia (HH), and hypobaric hypoxia + RSV (HH+RSV). Rats in the HH+RSV group received a continuous daily dose of 400 mg· kg^−1^ of RSV. Sterile, neutral 0.5% sodium carboxymethylcellulose solution served as the vehicle. All results were compared to Control or HH animals to evaluate improvements in the therapeutic potential of RSV under hypoxic conditions. Echocardiography and relevant indicators were monitored at the conclusion of study. Following the 2-weeks experimental period, the body weight and heart weight were measured. Heart tissue samples were collected for energy metabolism analysis, ferroptosis assessment, ATP measurement, and western blotting.

Cardiac hypertrophy was evaluated using the cardiac coefficient, calculated by the following formula: heart weight to body weight ratio.

### 2.3. Echocardiography measurements

At week 2, rats were anesthetized using 2% isoflurane. Subsequent echocardiography measurements were taken using a VINNO6LAB echocardiography system (Suzhou, China). M-mode recordings assessed left ventricular parameters, including cardiac output (CO), stroke volume (SV), left ventricular end-diastolic volume (LVEDV), left ventricular end-systolic volume (LVESV), and left ventricular posterior wall dimensions in diastole (LVPWd) and systole (LVPWs).

### 2.4. Euthanasia and tissue collection

After echocardiography, the rats were sacrificed by excess isoflurane (3-4%), then blood was taken from the abdominal aorta and tissue collected.

### 2.5. Transmission electron microscopy (TEM)

Left ventricular myocardial tissues were isolated and cut into small pieces (1 × 1 × 1 mm). The samples were fixed with 2.5% glutaraldehyde in phosphoric acid buffer for 2 h, followed by post-fixation in 1% osmium acid for 2 h. Ultrathin sections, approximately 50 nm thick, were prepared using an ultramicrotome. These slices were stained with 3% uranyl acetate and lead citrate for 15 min at room temperature. The mitochondrial structure was then observed using a TEM system (Tokyo, Japan).

### 2.6. Cell culture and experimental designs

H9c2 cells (from Pricella, Wuhan, China) were cultured in DMEM supplemented with 10% FBS and 1% penicillin-streptomycin at 37 °C with 5% CO_2_.

To stimulate cell damage, H9c2 cells were cultured in a hypoxia incubator with 0.5% O_2_ for varying durations: 0, 12, 24, 36, and 48 h. Finally, 48 h of hypoxia was confirmed as the optimal experimental duration.

In this study, H9c2 cells were divided into six groups: Control group, hypoxia group, hypoxia+Fer-1 group, hypoxia+WY14643 group, hypoxia+SIRT1^OE^ group and hypoxia+SIRT1^OE^+GW6471 group. H9c2 cells were subjected to 10 µmol/L Fer-1 for 6 h or 1 µmol/L WY14643 for 6 h, respectively. The culture medium was then refreshed, and the cells were treated in a hypoxia incubator (0.5% O₂) for an additional 48 h.

H9c2 cells underwent transfection with adenoviruses containing SIRT1 overexpression plasmids and Flag plasmids. The adenoviral titers used in this study were 3.5 × 10^10^ plaque-forming units (PFU)/mL, with a multiplicity of infection (MOI) of 50:1. Subsequent to adenoviral transfection for 10 h, the cells stably expressing SIRT1 were exposed to hypoxia for a duration of 48 h. Additionally, adenovirus transfection was co-cultured with GW6471 (0.25 µmol/L, 10 h) in hypoxia+SIRT1^OE^+GW6471 group. Then treated with hypoxia incubator (0.5% O_2_) for an additional 48 h. The cells were used for subsequent experiments after the intervention.

### 2.7. Measurement of cell viability

Normal H9c2 cells were cultivated into 96-well plate and exposed to Fer-1, WY14643, GW6471 for 6 h. The cells were then transferred to a hypoxia incubator with 0.5% O_2_ for 48 h. A CCK-8 assay was utilized for test H9c2 cell viability.

### 2.8. Measurement of ATP levels

After washing with precooled PBS, the treated cardiomyocytes were disrupted using a manual cell disruptor and quantified with the BCA Protein Assay Kit. The levels of ATP in myocardial tissue of rats or H9c2 cells were measured using the ATP Assay Kit.

### 2.9. Measurement of FFA, A-CoA, MDA, GSH and GSSG levels

FFA, A-CoA, MDA, GSH and GSSG levels in serum, myocardial tissue of rats, or H9c2 cells were detected using micro plate methods following the manufacturer’s instructions. The assay kits used have been described in the chemicals and reagents section.

### 2.10. The assay of ROS, mitochondrial membrane potential (Δψm), lipid peroxidation

The treated H9c2 cells were washed three times with PBS, followed by incubation with DCFH-DA at 37 ℃ 20 min. The nuclei were stained with hochest for 10 min in darkness, according to the manual. Similarly, TMRE staining was performed for 15 min to observe the mitochondrial membrane potential in treated H9c2 cells. For the observation of lipid peroxidation, H9c2 cells were treated with BODIPYTM 581/591 reagent working solution at 37 ℃ for 30 min in darkness. After washing, the stained cells were imaged under a fluorescent microscope.

To visualize ROS production in the heart tissue of the rats, the tissue slices were stained with 5 μM dihydroethidium and DAPI. Images were visualized and captured using a microscope.

### 2.11. Western blotting analysis

Total proteins were separated using RIPA Lysis Buffer containing protease inhibitors. Equal amounts of protein samples were loaded onto 4–12% SDS-PAGE gels. After the proteins were transferred to polyvinylidene fluoride membranes, the membranes were blocked with 5% milk for 2 h. The membranes were then incubated overnight at 4°C with specific primary antibodies for SIRT1, PPARα, CPT1, PDK4, GPX4, SLC7A11, and β-actin. A fluorescent secondary antibody was added to the membranes and incubated at room temperature for 1 h. The signals were visualized using Millipore ECL Western blotting detection reagents (Billerica, MA, USA) and the Tanon 5200 Gel Imaging System (Shanghai, China). The reaction products were densitometrically analyzed using an Odyssey infrared imaging system and ImageJ software. The ratio of the target protein was normalized to β-actin as the internal Control.

### 2.12. Statistical analysis

Statistical analysis was performed using SPSS 21.0 software. All data were expressed as the mean ± SD of at least three independent experiments. F and Kolmogorov–Smirnov tests were used to evaluate the homogeneity of variance and normality of distribution, respectively. Groups were compared using the Student’s t-test or one-way ANOVA with Tukey’s post hoc test, where appropriate. Values were considered to be statistically significant at *P < 0.05, **P < 0.01 vs. Control, #P < 0.05, ##P < 0.01 vs. HH or Hypoxia, &P < 0.05, &&P < 0.01 vs. Hypoxia+SIRT1^OE^.

## 3. Results

### 3.1. High-altitude hypoxia-induced cardiac injuries were accompanied by ferroptosis both *in vitro* and *in vivo*

To investigate the effect of high-altitude on rat heart structure and function, echocardiography was conducted. The results revealed significant decreases (P < 0.05; Fig. 1A-D) in the levels of LVEDV, LVESV, SV and CO in the HH group compared to the Control group. Additionally, LVPWd and LVPWs showed significant increases (P < 0.05; Fig. 1E) in the HH group. Furthermore, the heart weight-to-body ratios increased (P < 0.05; Fig. 1F) in the HH group, indicating the successful construction of a cardiac injury model in high-altitude hypoxia. And in H9c2 cells, as hypoxia persisted, cell viability progressively decreased (P < 0.05; Fig. 1J). Moreover, it revealed a significant increase in ROS levels (P < 0.05; Fig. 1G, I) and MDA levels (P < 0.05; Fig. 1K, L) in myocardial cells exposed to high-altitude hypoxia both *in vitro* and *in vivo*. Given that elevated ROS can induce ferroptosis^[26]^, we investigated other indicators associated with ferroptosis. The results indicated a marked decrease in GSH/GSSG contents (P < 0.05; Fig 1M), suggesting the conversion of glutathione from its reduced to oxidized form^[27]^. Additionally, the GSH/GSSG contents gradually decreased with prolonged exposure to hypoxia time in H9c2 cells (P < 0.05; Fig. 1N). Furthermore, reduced protein levels of GPX4, a hallmark protein of ferroptosis^[28]^, were observed both in rat myocardial tissue (P < 0.05; Fig. 1O) and H9c2 cells (P < 0.05; Fig. 1P) in response to high-altitude hypoxia exposure. These findings collectively indicate an increase in ferroptosis both *in vitro* and *in vivo* hypoxia exposure.

**Figure 1.**
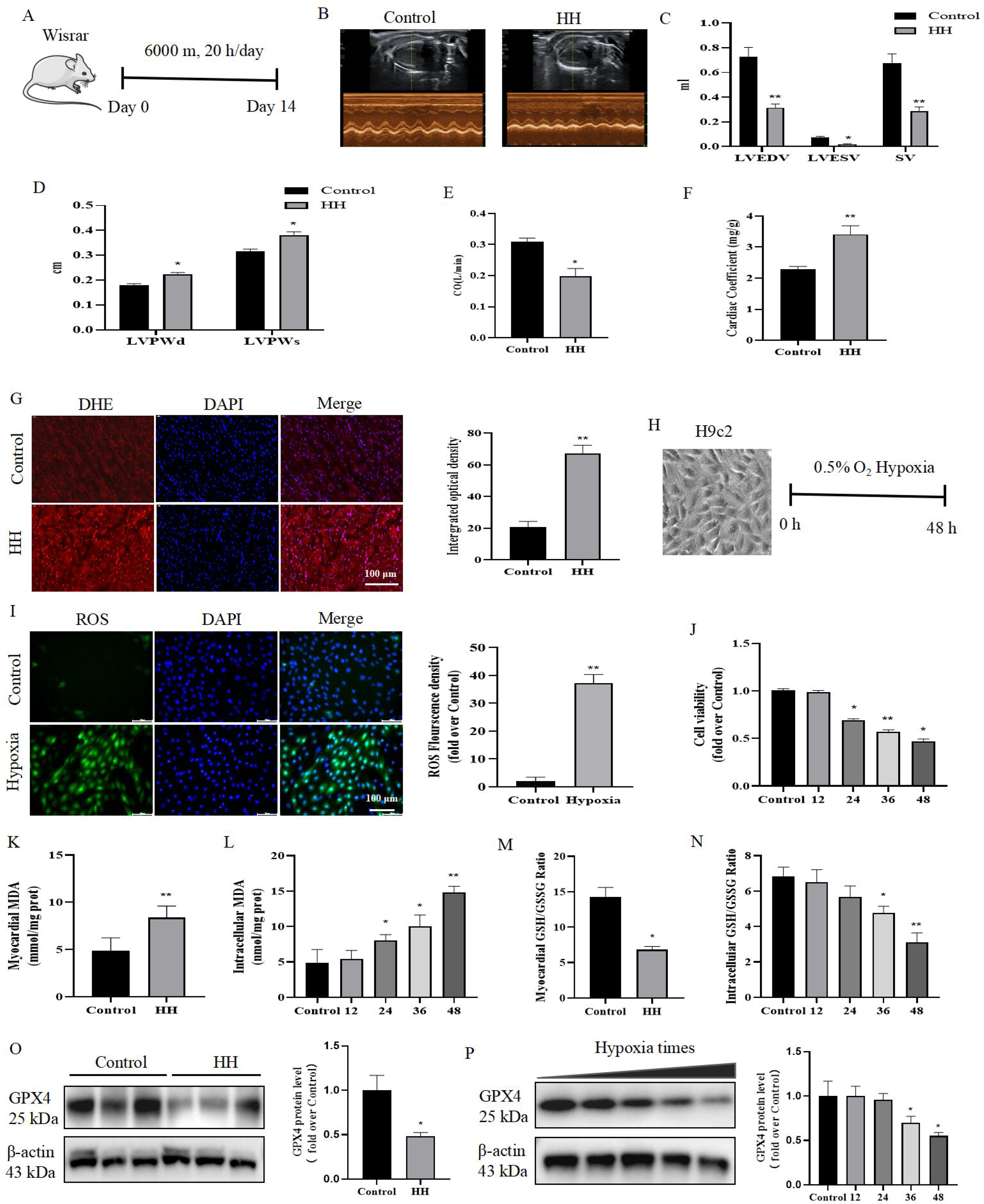
**High-altitude hypoxia-induced cardiac injuries were accompanied by ferroptosis both *in vitro* and *in vivo.*** (A) Animal study design. (B) Echocardiography in rats. (C-E) HH-exposed rats showed decreased levels of LVEDV, LVESV, SV, and CO, and increased levels of LVPWd, LVPWs compared with Control. (F) HH-exposed rats exhibited an elevated cardiac coefficient compared with Control. (G) DHE staining of rat myocardial tissue (40×) after HH induction and quantitative analysis of fluorescence. (H) Celluar study design. (I) DCFH-DA staining and quantitative analysis of fluorescence in hypoxia-induced H9c2 cells. (J) H9c2 cell viability after different durations of hypoxia exposure, assessed using a CCK-8 assay. (K) Levels of MDA in myocardial tissue. (L) Levels of MDA in H9c2 cells. (M) Levels of GSH/GSSG in myocardial tissue. (N) Levels of GSH/GSSG in H9c2 cells. (O) Western blotting was performed on rat myocardial tissue extracts to determine GPX4 levels, and grayscale values were analyzed, with β-actin serving as a protein loading control. (P) H9c2 cardiomyocyte extracts were subjected to western blotting to measure GPX4 levels, and grayscale values were analyzed, with β-actin serving as a protein loading control. Statistical significance was indicated as *P < 0.05, **P < 0.01 vs. Control.

### 3.2. Inhibition of ferroptosis alleviated hypoxia-induced cardiomyocyte injury

We further explored the role of ferroptosis in cardiomyocyte injury caused by hypoxia exposure (Fig. 2A). Fer-1 is recognized as a potent inhibitor of ferroptosis, as it promotes cystine import and GSH production^[29]^. Compared to the hypoxia group, western blotting results indicated that Fer-1 treatment significantly increased the protein expression of GPX4 and SLC7A11 (P < 0.05; Fig. 2B). Additionally, pretreatment of Fer-1 resulted in a significant reduction in ROS levels in hypoxia-induced H9c2 cells (P < 0.05; Fig. 2C). Furthermore, the increased MDA levels (P < 0.05; Fig. 2D) and decreased GSH/GSSG ratio (P < 0.05; Fig. 2E) observed in the hypoxia group were restored by Fer-1 treatment. Notably, Fer-1 pretreatment led to a significant increase in hypoxia-induced cell viability (P < 0.05; Fig. 2F). Collectively, these findings suggest that the ferroptosis specific inhibitor Fer-1 can reverse ferroptosis-related changes in hypoxia-induced H9c2 cells. This further supports the notion that ferroptosis plays a role in cardiac injury during hypoxic conditions.

**Figure 2.**
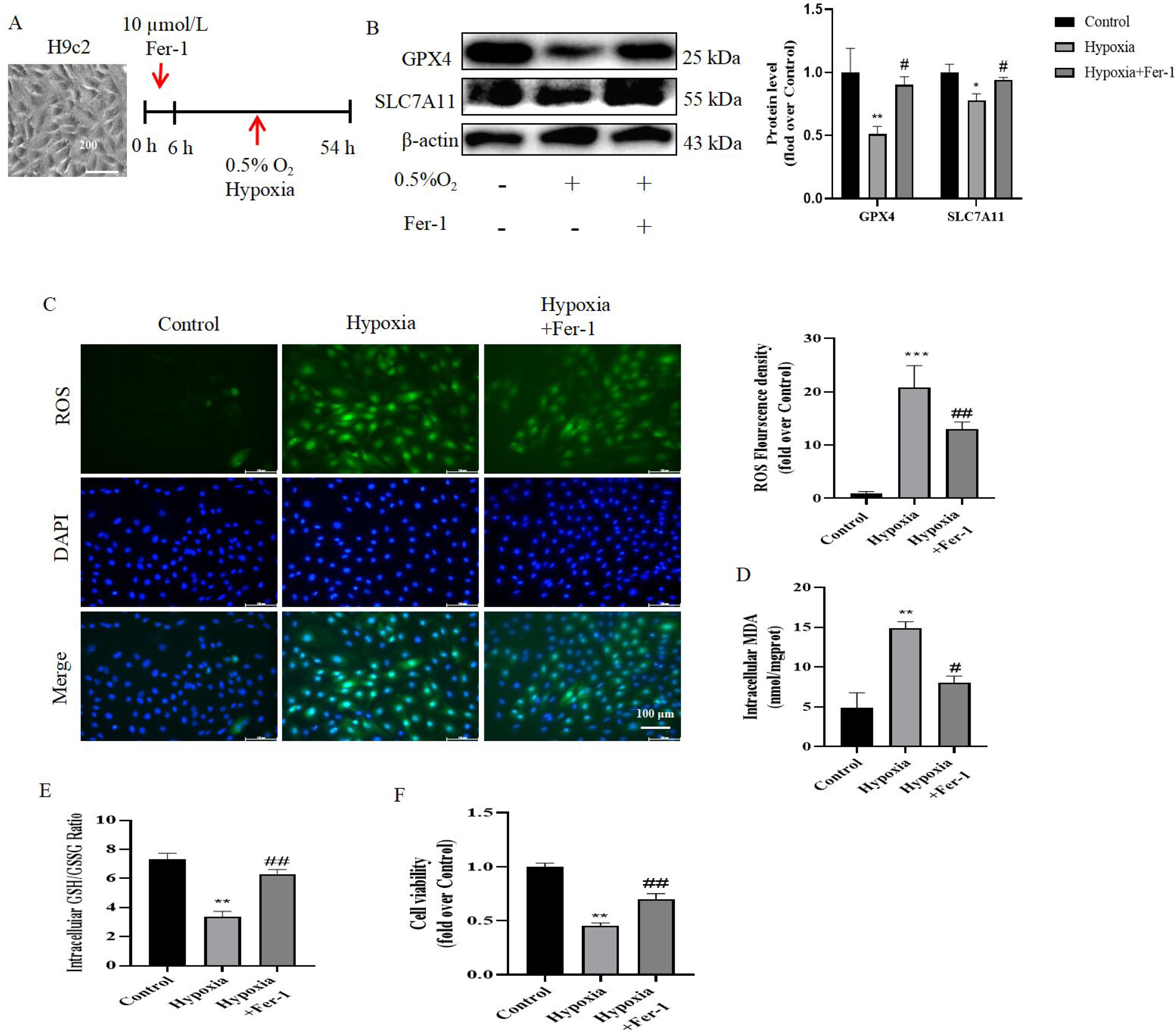
Inhibition of ferroptosis alleviated hypoxia-induced cardiomyocyte injury. (A) Celluar study design. (B) H9c2 cardiomyocyte extracts were subjected to western blotting to measure GPX4 and SLC7A11 levels, and grayscale values were analyzed, with β-actin serving as a protein loading control. (C) After Fer-1 pretreatment, representative images of DCFH-DA staining and quantitative analysis of fluorescence in H9c2 cells. Scale bar = 100 μm. (D) Intracellular MDA levels in H9c2 cells after Fer-1 pretreatment. (E) Intracellular GSH/GSSG levels in H9c2 cells after Fer-1 pretreatment. (F) CCK-8 assays were used to assess the viability of H9c2 cells after Fer-1 pretreatment. Statistical significance was indicated as *P < 0.05, **P < 0.01 vs. Control, #P < 0.05, ##P < 0.01 vs. Hypoxia.

### 3.3. Changes in mitochondrial structure and function, and FAO in high-altitude hypoxia-induced myocardial injury both *in vitro* and *in vivo*

Subsequently, we investigated the left ventricular myocardial mitochondrial structure in rats. Transmission electron microscopy indicated that the mitochondrial structure in the HH group displayed blurred, swollen, and fragmented cristae (Fig. 3A). A detailed analysis of the number of mitochondria per unit area and the mean mitochondrial size showed an increase in the number of mitochondria per unit area in the HH group (P < 0.05; Fig. 3B), while the mean mitochondrial size decreased (P < 0.05; Fig. 3C) compared to those in the Control group. The mitochondrial membrane potential is essential for maintaining mitochondria function^[30]^. TMRE staining showed lower levels of red fluorescence intensity as hypoxia persisted compared with the control group (P < 0.05; Fig. 3D). Mitochondria serve as the primary energy suppliers for myocardial function^[31]^. Analysis of high-altitude hypoxia-induced rat myocardial tissue revealed an increase in FFA and A-CoA levels (P < 0.05; Fig. 3E, G), and a simultaneous decrease in ATP levels (P < 0.05; Fig. 3I) compared to the Control group. Similarly, when H9c2 cells were exposed to hypoxia, FFA and A-CoA levels gradually increased (P < 0.05; Fig. 3F, H) and ATP levels gradually decreased (P < 0.05; Fig. 3J). PPARα plays a crucial role in cellular fatty acid uptake, esterification, and trafficking, while also regulating genes involved in lipoprotein metabolism^[32]^. Notably, reduced protein levels of PPARα were observed in both rat myocardial tissue (P < 0.05; Fig. 3K) and H9c2 cells (P < 0.05; Fig. 3L) in response to hypoxia exposure. These results indicate that FAO is reduced in both *in vitr*o and *in vivo* hypoxia exposure.

**Figure 3.**
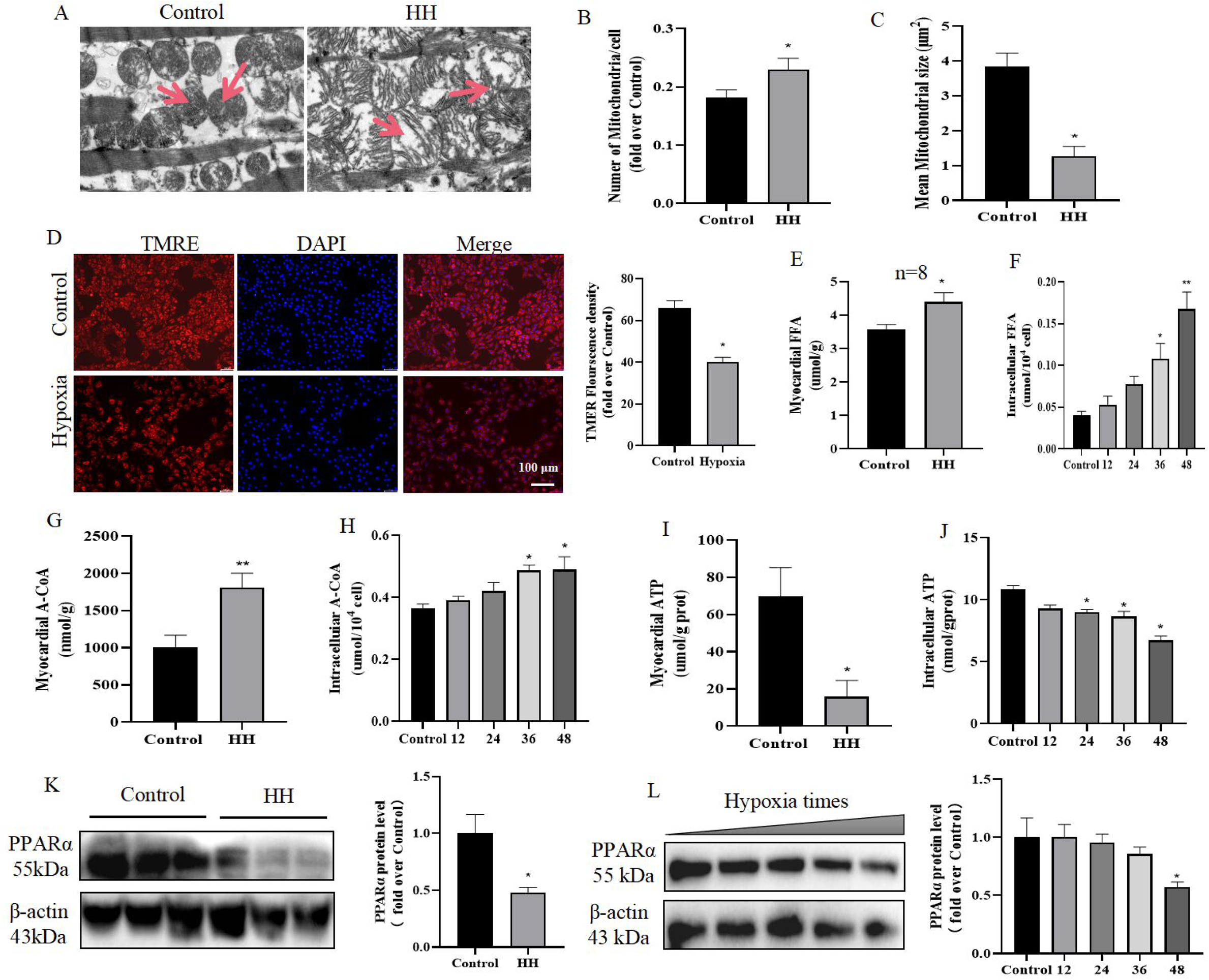
Changes in mitochondrial structure and function, and FAO in high-altitude hypoxia-induced myocardial injury both *in vitro* and *in vivo.* (A) Transmission electron microscopy was utilized to visualize the left ventricular myocardium ultrastructure in rats (8,000×). Scale bar = 1 μm. (B, C) A detailed analysis was conducted to determine the number of mitochondria per unit area and the average mitochondrial size. (D) After different durations of hypoxia exposure, representative images of TMRE staining and quantitative analysis of fluorescence in H9c2 cells. Scale bar = 100 μm. Levels of myocardial FFA (E), A-CoA (G), and ATP (I) in HH-exposed rats. Intracellular levels of FFA (F), A-CoA (H), and ATP (J) in H9c2 cells after hypoxia exposure. (K) Western blotting was performed on rat myocardial tissue extracts to determine PPARα levels, and grayscale values were analyzed, with β-actin serving as a protein loading control. (L) H9c2 cardiomyocyte extracts were subjected to western blotting to measure PPARα levels, and grayscale values were analyzed, again using β-actin as a protein loading control. Statistical significance was indicated as *P < 0.05, **P < 0.01 vs. Control.

### 3.4. Activation of PPARα improves FAO and suppresses hypoxia-induced ferroptosis

To further validate the role of PPARα in hypoxia-induced H9c2 cells, we activated PPARα using WY14643 (Fig. 4A), a specific agonist^[33]^. In the hypoxia group, the expression of the FAO-associated proteins PPARα, PDK4 and CPT1 was lower compared to the Control group. Following simultaneous treatment with WY14643, the expression levels of these proteins gradually increased (P < 0.05; Fig. 4B). Moreover, after treatment of H9c2 cells with WY14643, intracellular levels of FFA and A-CoA decreased (P < 0.05; Fig. 4C, D) while intracellular mitochondrial ATP concentration increased (P < 0.05; Fig. 4E). Furthermore, TMRE staining showed higher mitochondrial membrane potential with WY14643 treatment (P < 0.05; Fig. 4G, I). To verify whether PPARα is involved in ferroptosis caused by hypoxia, we assessed GPX4 and SLC7A11 protein expression. The results showed that WY14643 could improve the protein expression of GPX4 and SLC7A11 (P < 0.05; Fig. 4B) in hypoxia-induced H9c2 cells. Meanwhile, the PPARα agonist WY14643 increased GSH/GSSG ratio (P < 0.05; Fig. 4F) in hypoxia-induced H9c2 cells. Additionally, TMRE staining showed lower ROS levels (P < 0.05; Fig. 4H, J) with WY14643 treatment. Additionally, in hypoxia-treated H9c2 cells, WY14643 activated PPARα, significantly increasing cell viability (P < 0.05; Fig. 4K). Collectively, these results indicate that when exposed to hypoxia, FAO decreased, and ferroptosis became more pronounced, whereas this trend was improved by WY14643.

**Figure 4.**
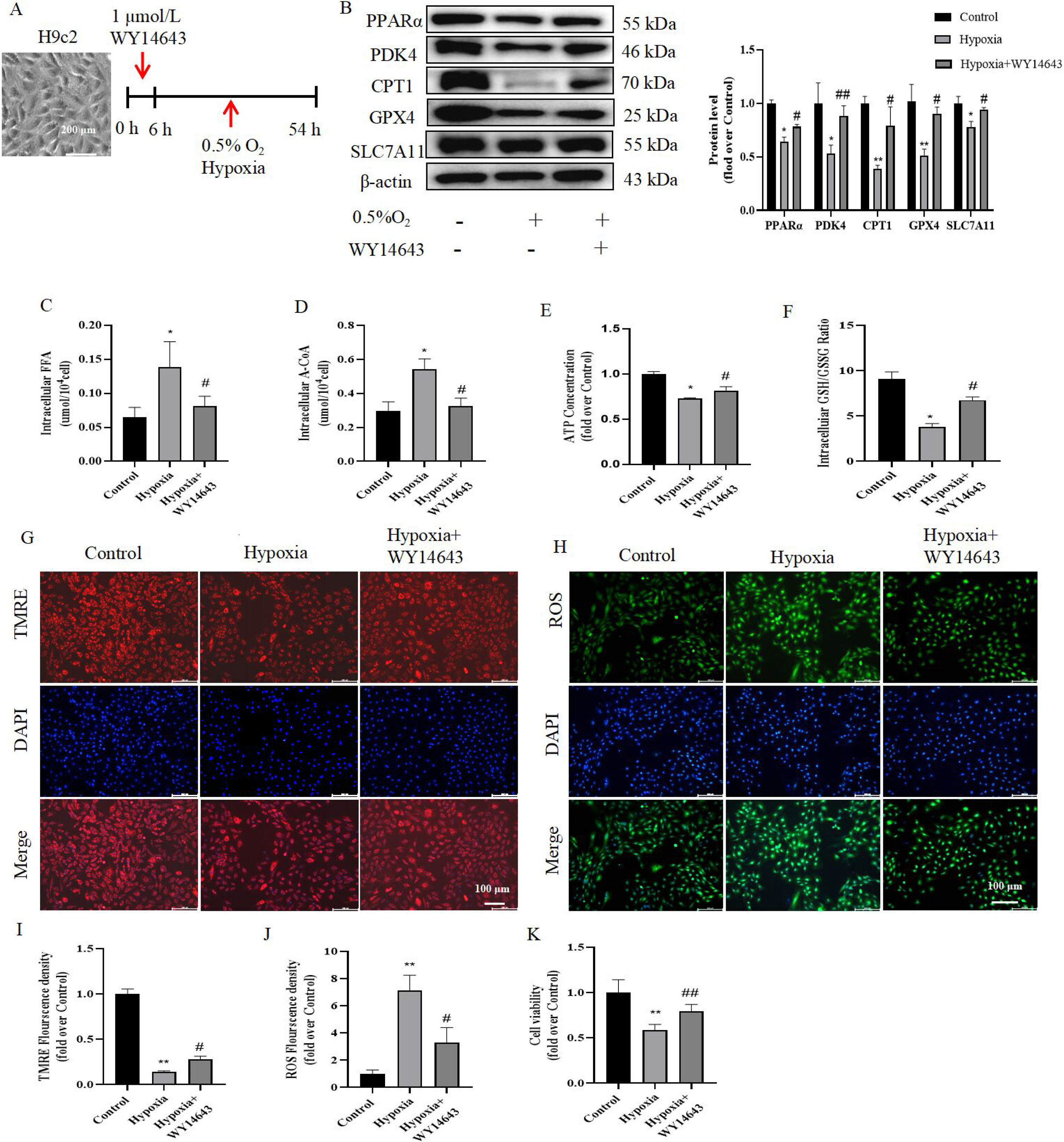
Activation of PPARα improves FAO and suppresses hypoxia-induced ferroptosis. (A) Celluar study design. (B) After PPARα activation, western blotting was performed on H9c2 cells to determine PDK4, CPT1, GPX4, and SLC7A11 levels, and grayscale values were analyzed, with β-actin serving as a protein loading control. Intracellular levels of FFA (C) and A-CoA (D) after treatment of H9c2 cells with WY14643. (E) Intracellular ATP levels in H9c2 cells after PPARα activation. (F) Intracellular GSH/GSSG levels in H9c2 cells after PPARα activation. (G) After PPARα activation, representative images of TMRE staining and quantitative analysis of fluorescence (I) in H9c2 cells. Scale bar = 100 μm. (H) After PPARα activation, representative images of DCFH-DA staining and quantitative analysis of fluorescence (J) in H9c2 cells. Scale bar = 100 μm. (K) CCK-8 assays were employed to assess the viability of H9c2 cells after PPARα activation. Statistical significance was indicated as *P < 0.05, **P < 0.01 vs. Control, #P < 0.05, ##P < 0.01 vs. Hypoxia.

### 3.5. Overexpression of SIRT1 improves FAO and suppresses hypoxia-induced ferroptosis

To explore the regulatory mechanism of PPARα on cardiac energy metabolism under hypoxia, we further investigated the role of SIRT1, the upstream regulator of PPARα (Fig. 5A). Monounsaturated fatty acids enhance PGC1α/PPARα signaling and promote oxidative metabolism in cells and animal models in a SIRT1-dependent manner^[34]^. Overexpression of the FLAG-tagged fusion protein SIRT1, using adenoviral vectors, elevated the protein expression of PGC1α, PPARα, PDK4 and CPT1 (P < 0.05; Fig. 5B) compared to those in the hypoxic group. Futhermore, after overexpression of SIRT1 in H9c2, the intracellular FFA and A-CoA levels decresed (P < 0.05; Fig. 5C, D) and ATP levels increased (P < 0.05; Fig. 5E) compared with levels in the hypoxic group. And TMRE staining showed that overexpression of SIRT1 significantly increased mitochondrial membrane potential (P < 0.05; Fig. 5G, I). These results indicate that upregulation of SIRT1 can improve hypoxia-induced decrease in FAO in H9c2 cardiomyocytes. To further verify the effect of SIRT1 regulation on PPARα in enhancing the efficacy of fatty acid oxidation (FAO) in mitigating ferroptosis, we examined GPX4 and SLC7A11 protein expression. The results showed that overexpression of SIRT1 increased the expression of GPX4 and SLC7A11 in hypoxia-induced H9c2 cardiomyocytes (P < 0.05; Fig. 5B). Moreover, overexpression of SIRT1 significantly increased the level of GSH/GSSG ratio (P < 0.05; Fig. 5F) and reduced ROS (P < 0.05; Fig. 5H, J) compared with the hypoxia group. Additionally, Overexpression of SIRT1 also significantly enhanced cell viability (P < 0.05; Fig. 5K). These findings suggest that the overexpression of SIRT1 can upregulate PPARα and PGC1α, thereby enhancing FAO and reducing ferroptosis induced by hypoxia in H9c2 cells.

**Figure 5.**
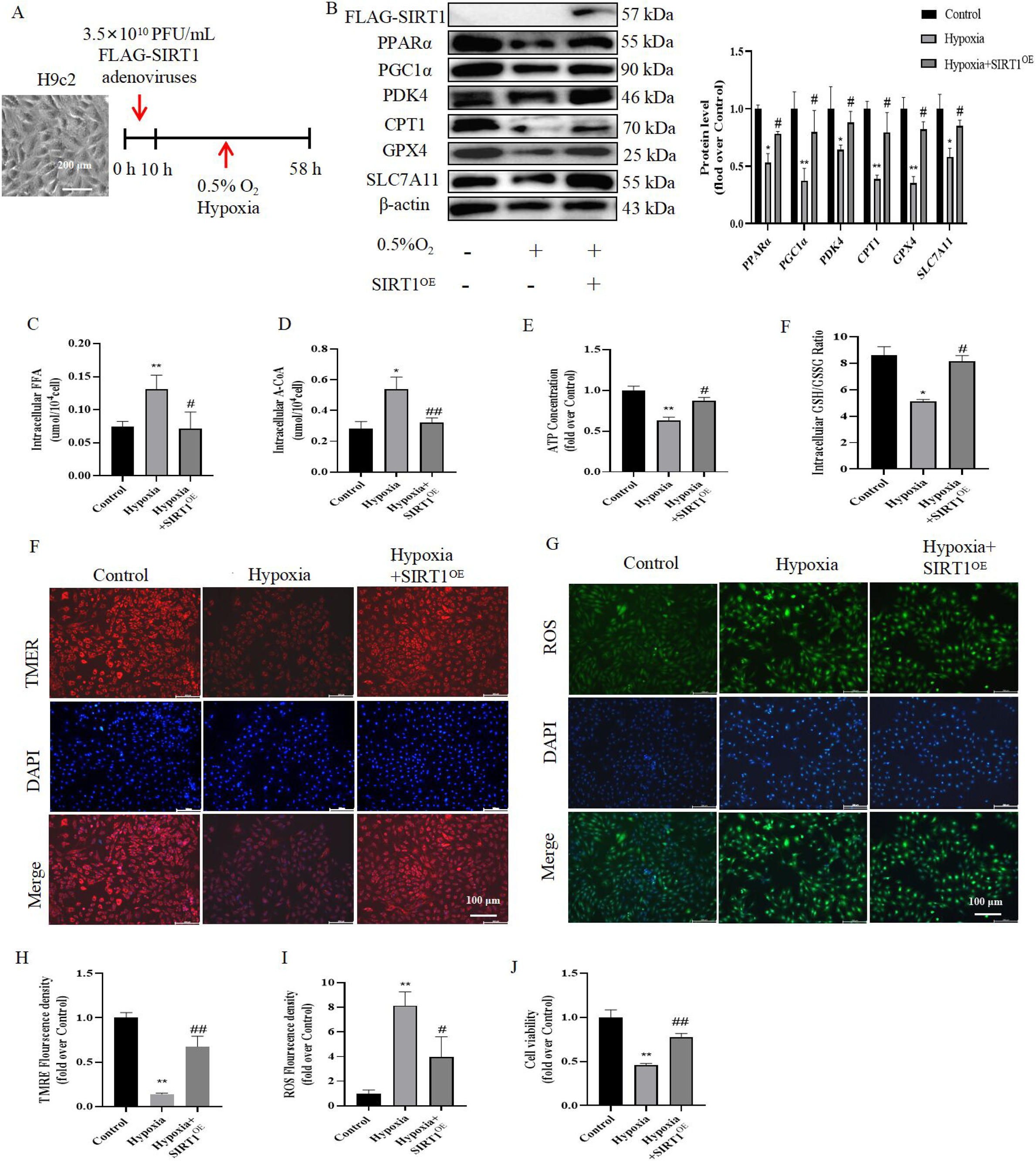
Overexpression of SIRT1 improves FAO and suppresses hypoxia-induced ferroptosis. (A) Celluar study design. (B) After SIRT1 overexpression, western blotting was performed on H9c2 cells to determine PGC1α, PPARα, PDK4, CPT1, GPX4, and SLC7A11 levels, and grayscale values were analyzed, with β-actin serving as a protein loading control. Intracellular levels of FFA (C) and A-CoA (D) after SIRT1 overexpression in H9c2 cells. (E) Intracellular ATP levels in H9c2 cells after SIRT1 overexpression. (F) Intracellular GSH/GSSG levels in H9c2 cells after SIRT1 overexpression. (G) Representative images of TMRE staining and quantitative analysis of fluorescence (I) in H9c2 cells after SIRT1 overexpression. Scale bar = 100 μm. (H) Representative images of DCFH-DA staining and quantitative analysis of fluorescence (J) in H9c2 cells after SIRT1 overexpression. Scale bar = 100 μm. (K) CCK-8 assays were employed to assess the viability of H9c2 cells after SIRT1 overexpression. Statistical significance was indicated as *P < 0.05, **P < 0.01 vs. Control, #P < 0.05, ##P < 0.01 vs. Hypoxia.

### 3.6. The regulation of SIRT1 on FAO and ferroptosis under hypoxic conditions requires the involvement of PPARα

PPARα is essential for the regulation of SIRT1 on FAO and ferroptosis under hypoxic conditions. To delve into the relationship betweenSIRT1, PPARα, FAO and ferroptosis, we utilized the PPARα inhibitor GW6471 while overexpressing SIRT1 (Fig. 6A). As depicted in Fig. 6B, overexpression of SIRT1 led to increased protein expression levels of PGC1α, PPARα, PDK4, CPT1, GPX4 and SLC7A11 in hypoxia-treated H9c2 cells. However, the application of the PPARα inhibitor GW6471 negated the effects of SIRT1 overexpression on these protein levels in hypoxia-treated H9c2 cells (P < 0.05, Fig. 6B). Moreover, GW6471 treatment, targeting PPARα, obstructed the influence of SIRT1 overexpression on the intracellular levels of FFA, A-CoA and ATP levels in hypoxia-induced H9c2 cells (P < 0.05, Fig. 6C-E). And GW6471 treatment resulted in a marked decrease in mitochondrial membrane potential (P < 0.05, Fig. 6G, I). Subsequent analysis of the ROS levels and GSH/GSSG ratio revealed that GW6471 significantly diminished the GSH/GSSG ratio (P < 0.05, Fig. 6F) and elevated ROS levels (P < 0.05, Fig. 6H, J) in hypoxia-treated H9c2 overexpressing SIRT1. Additionally, GW6471 treatment resulted in a marked decrease in cell viability (P < 0.05, Fig. 6K), demonstrating that inhibiting PPARα can significantly counteract the protective effects of SIRT1 overexpression on cardiomyocytes FAO, ferroptosis, and myocardial damage. These outcomes suggest that the SIRT1-PPARα-GPX4 signaling pathwayis crucial in modulating hypoxia-induced FAO and ferroptosis.

**Figure 6.**
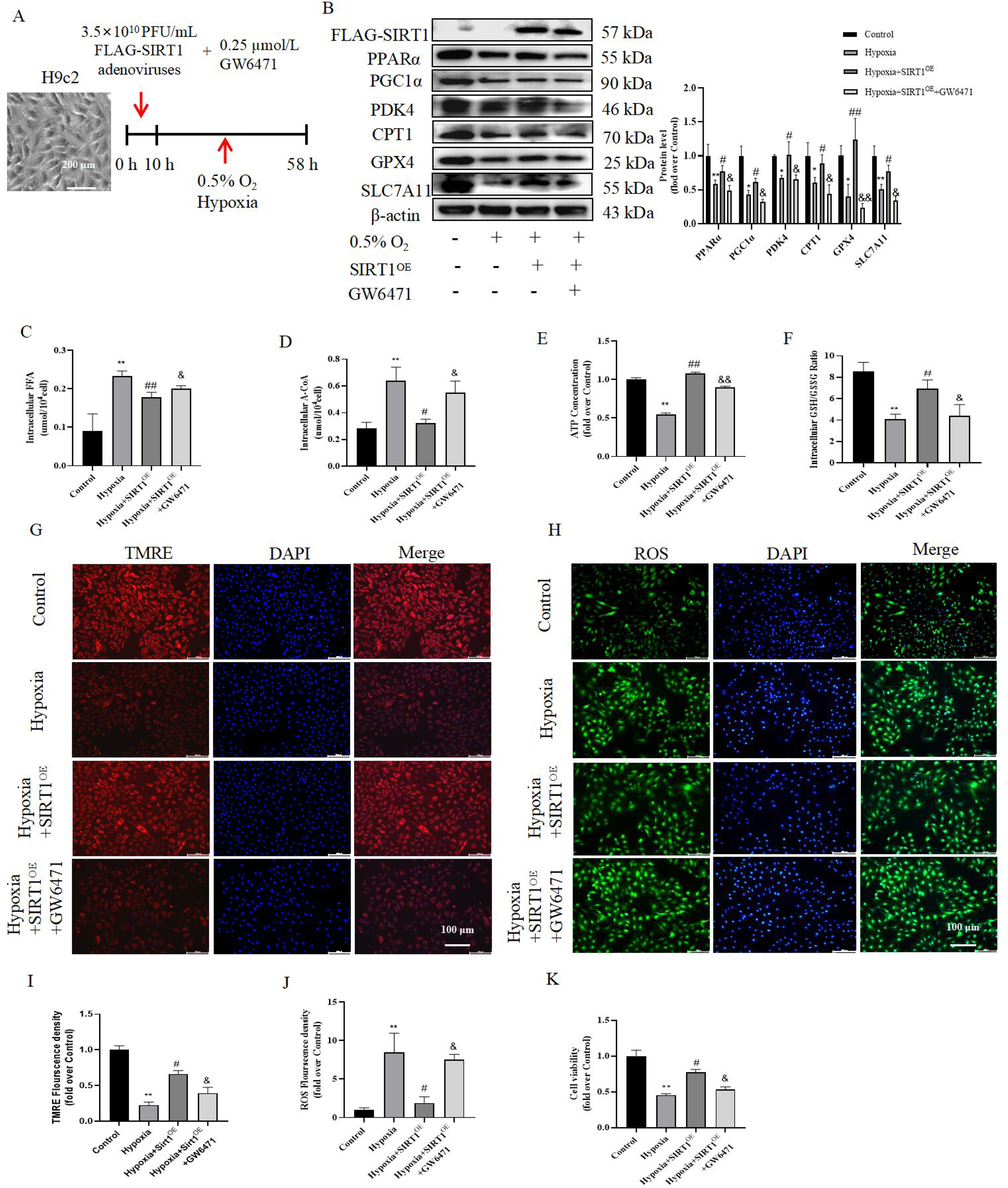
The regulation of SIRT1 on FAO and ferroptosis under hypoxic conditions requires the involvement of PPARα SIRT1. (A) Celluar study design. (B) After overexpression of SIRT1, PPARα was inhibited by GW6471 in H9c2 cells. Western blotting was conducted to measure the levels of PGC1α, PPARα, PDK4, CPT1, GPX4, and SLC7A11 in H9c2 cardiomyocytes, and grayscale values were analyzed. β-actin was used as a protein loading control. Intracellular levels of FFA (C), A-CoA (D), ATP (E), and GSH/GSSG (F) in H9c2 cells after SIRT1 overexpression and PPARα inhibition. (G) Representative images of TMRE staining and quantitative analysis of fluorescence (I) in H9c2 cells after SIRT1 overexpression and PPARα inhibition. Scale bar = 100 μm. (H) Representative images of DCFH-DA staining and quantitative analysis of fluorescence (J) in H9c2 cells after SIRT1 overexpression and PPARα inhibition. Scale bar = 100 μm. (K) CCK-8 assays were employed to assess the viability of H9c2 cells after SIRT1 overexpression and PPARα inhibition. Statistical significance was indicated as *P < 0.05, **P < 0.01 vs. Control, #P < 0.05, ##P < 0.01 vs. Hypoxia, &P < 0.05, &&P < 0.01 vs. Hypoxia+SIRT1^OE^.

### 3.7. Resveratrol improves FAO and ferroptosis by regulating SIRT1-PPARα-GPX4 pathway in hypobaric hypoxia-induced cardiac injury

Resveratrol,a natural activator of SIRT1^[35]^. It was evaluated for its potential to alleviate myocardial injury induced by high altitude, focusing on improvements in myocardial energy metabolism and ferroptosis. Echocardiography results indicated that resveratrol administration significantly increased the levels of LVEDV, LVESV, SV and CO (P < 0.05; Fig. 7A-D) and decreased LVPWd, LVPWs and cardiac coefficient (P < 0.05; Fig. 7E, F) compared with the HH group. Moreover, resveratrol treatment significantly reduced the number of mitochondria per unit area and increased the mean mitochondrial size in the left ventricular of HH-induced rats (P < 0.05; Fig. 7G-I) compared to the HH group. Further analysis showed that the levels of FFA and A-CoA in HH group were increased, while levels of ATP decreased compared with the control group, resveratrol administration significantly reversed these changes in FAO indicators (P < 0.05; Fig. 7J-L). Additionally, DHE stainingshowed that resveratrol treatment significantly blunted the increase in ROS levels induced by HH exposure (P < 0.05; Fig. 7M, O), and significantly increased the myocardial GSH/GSSG ratio compared with the HH group (P < 0.05; Fig. 7P). More importantly, treatment with resveratrol also significantly improved the levels of SIRT1, PGC1α, PPARα, PDK4, CPT1, GPX4 and SLC7A11 protein in HH-induced rat hearts (P < 0.05; Fig. 7N, Q). These findings suggested that SIRT1’s natural agonist, resveratrol, regulates myocardial energy metabolism and ferroptosis through the SIRT1-PPARα-GPX4 axis, thereby improving cardiac function following exposure to high-altitude hypoxia.

**Figure 7.**
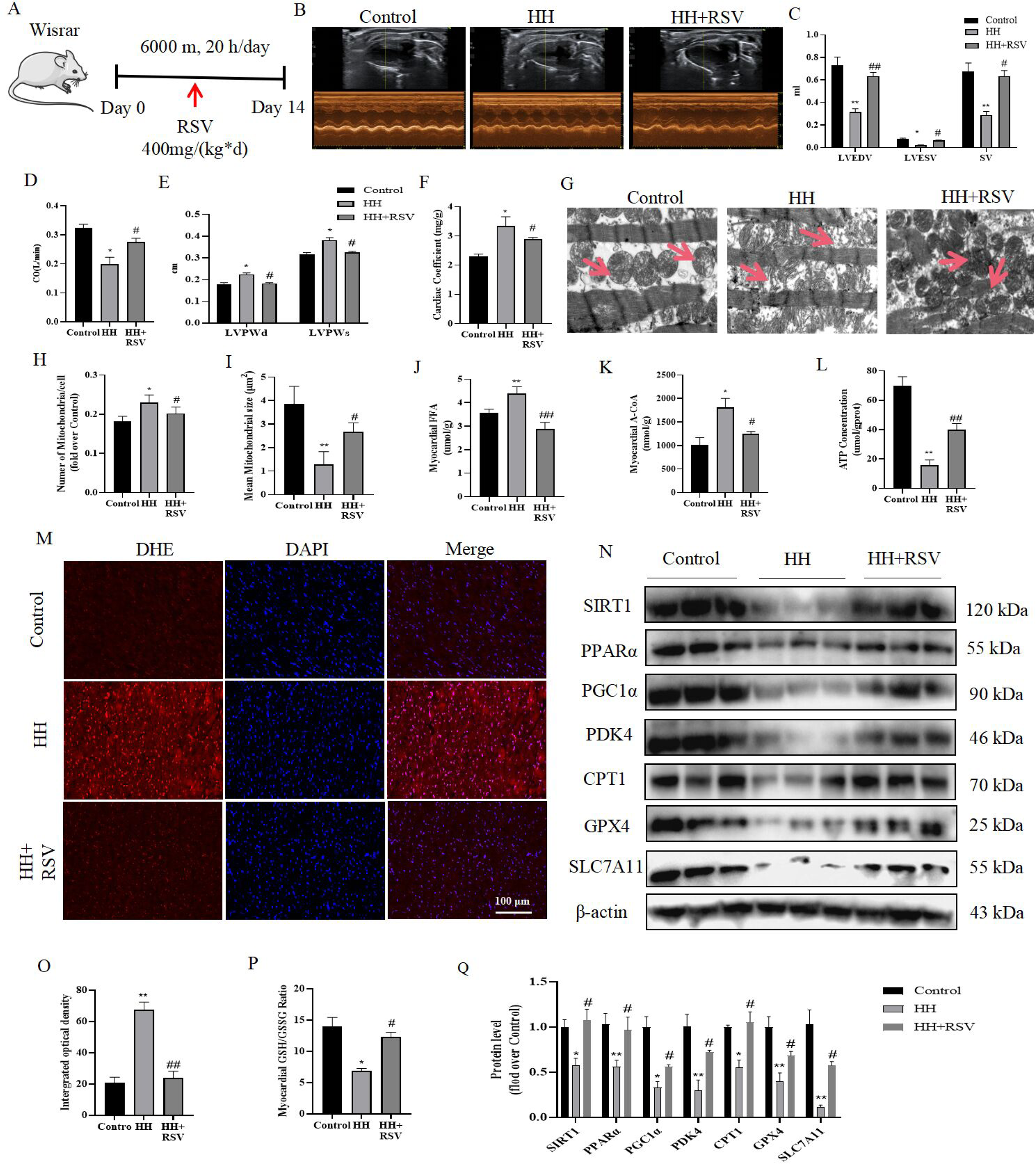
Resveratrol improves FAO and ferroptosis by regulating SIRT1-PPARα-GPX4 pathway in hypobaric hypoxia-induced cardiac injury. (A) Animal study design. (B) Echocardiography in rats. (C-E) RSV decreased LVPWd and LVPWs levels and increased LVEDV, LVESV, SV, and CO levels of the heart in the HH test. (F) RSV decreased the cardiac coefficient compared with the HH group. (G) Transmission electron microscopy was utilized to visualize the left ventricular myocardium ultrastructure in rats (8,000×). Scale bar = 1 μm. (H, I) A detailed analysis was conducted to determine the number of mitochondria per unit area and the average mitochondrial size. Myocardial FFA (J), A-CoA (K), and ATP (L) levels in rats after treatment with RSV. (M) DHE staining of rat myocardial tissue (40×) after RSV treatment and quantitative analysis of fluorescence (O). (N) Western blotting was performed on rat myocardial tissue extracts to determine SIRT1, PGC1α, PPARα, PDK4, CPT1, GPX4, and SLC7A11 levels, and grayscale values were analyzed (Q), with β-actin serving as a protein loading control. (P) Myocardial GSH/GSSG levels in rats after treatment with RSV. Statistical significance was indicated as *P < 0.05, **P < 0.01 vs. Control, #P < 0.05, ##P < 0.01 vs. HH.

Altogether, the data show that PPARα is regulated by SIRT1 in hypoxia-induced cardiac injury and is involved in myocardial energy metabolism and ferroptosis. Upregulation of PPARα can ameliorate hypoxia-induced cardiac injury by improving FAO and ferroptosis through CPT1/PDK4-GPX4/SLC7A11 pathway. (Figure 8).

**Figure 8.**
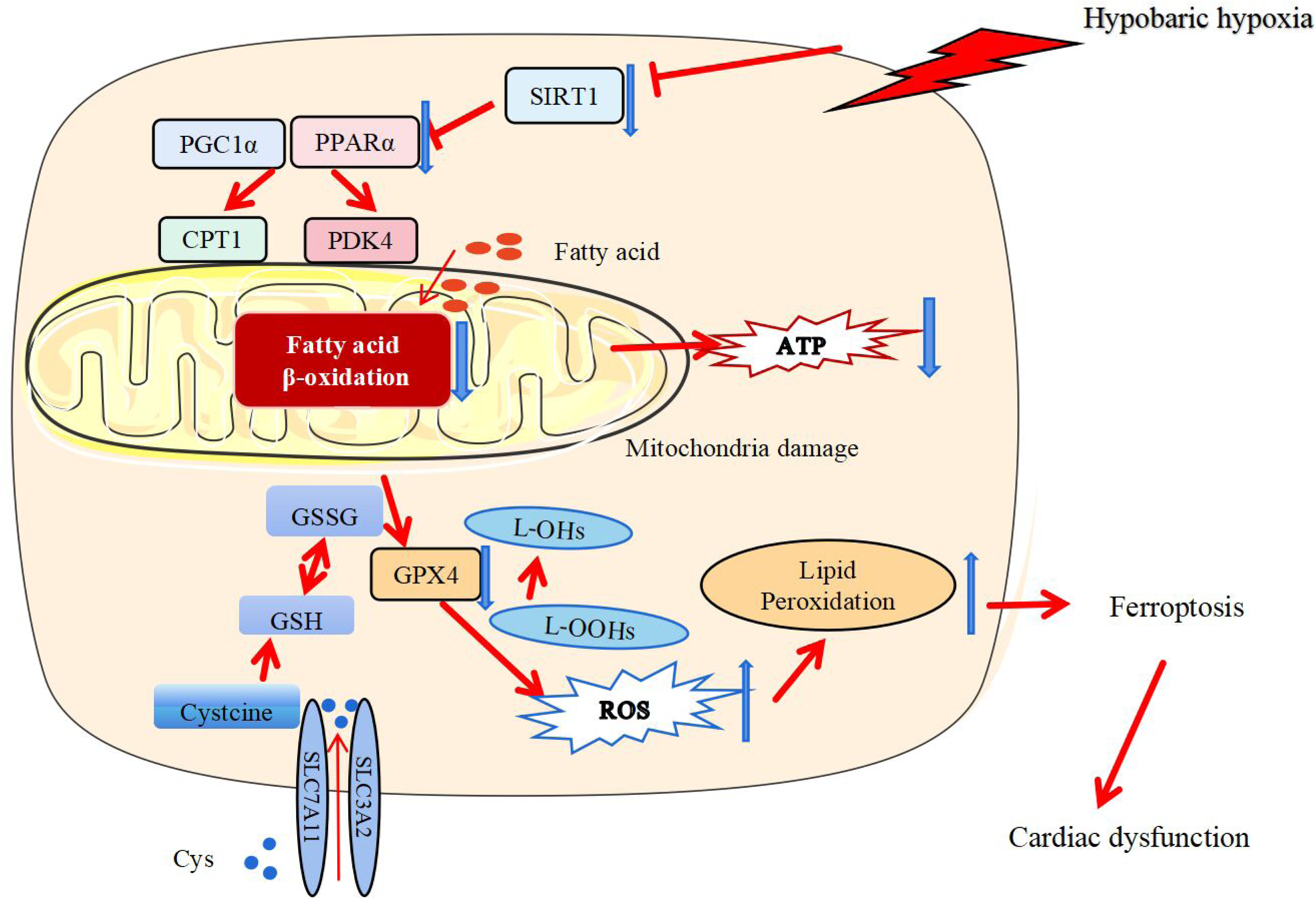
**Under conditions of HH exposure, the downregulation of SIRT1 triggers FAO and ferroptosis through the PPARα-PDK4/CPT1-GPX4/SLC7A11 pathway. This results in mitochondrial dysfunction, ultimately contributing to the development of cardiac impairment, as depicted in the pathway map.**

## 4. Discussion

The high-altitude environment is characterized by hypobaric hypoxia, low temperatures, and low humidity^[36]^. It has been reported that high altitude could induce irreversible or reversible damage to high oxygen and energy demanding tissues, such as the brain, heart, liver, over time^[37–39]^. Due to its high oxygen demand, the heart is particularly sensitive to hypoxia. In our study, rats were exposed to a hypobaric chamber at an altitude of 6000 m for two weeks. Echocardiography revealed significant changes in the heart structure, characterized by increased left ventricular posterior wall dimension and decreased cardiac output. The increase in cardiac coefficient indicated the development of cardiac hypertrophy in rats induced by HH. Similarly, H9c2 cells exposed to 0.5% O_2_ in a hypoxia incubator for 48 h exhibited significantly reduced cell viability. Fluorescent staining of rat myocardial tissue also indicated increased ROS levels induced by hypoxia. Since ROS serves as a key marker of ferroptosis, we speculate that ferroptosis occurs in the myocardium under hypoxia. Initially regarded as a catastrophic breakdown of cellular functions induced by chemical and physical stressors, ferroptosis is now recognized as a convergence of disrupted metabolic pathways and impaired cellular defense mechanisms^[24]^. Glutathione can cycle between reduced (GSH) and oxidized (GSSG) states, enabling this metabolite to participate in redox biochemical reactions. Glutathione peroxidases (GPXs) are evolutionarily highly conserved enzymes that use GSH as a cofactor to reduce peroxides (e.g., R–OOH) to their corresponding alcohols (R–OH), thereby limiting the transition metal-dependent formation of toxic radicals (e.g., R-O•)^[6]^. Our study found decreased expression levels of GPX4, a key regulator of ferroptosis, and a decrease in the GSH/GSSG ratio both *in vivo* and *in vitro* under hypoxic conditions.

To investigate the involvement of ferroptosis in cardiac injury under hypoxia, we examined the effects of Fer-1, a ferroptosis inhibitor. Fer-1 has been reported to effectively inhibit ROS overproduction and lipid peroxidation, which ultimately blocked ferroptosis process^[29]^. Fer-1 treatment restored GPX4 and SLC7A11 expression levels, reduced ROS and MDA levels, and increased the GSH/GSSG ratio compared to hypoxia conditions in H9c2 cells. Moreover, Fer-1 pretreatment increased cell viability, further suggesting that ferroptosis may contribute to hypoxia-induced cardiac injury.

Subsequently, we observed swollen and fragmented mitochondria by electron microscopy in rat myocardial tissue. Similarly, we found a significant decrease mitochondrial membrane potential in hypoxia-induced H9c2 cells. Mitochondria are complex organelle that play essential roles in energy transduction, ATP production, and various cellular signaling events^[40]^. The high-capacity mitochondrial system in the heart is dynamically regulated to generate and consume enormous amounts of ATP, supporting its constant pumping function amidst varying energy demands^[41]^. Mitochondrial damage often correlates with alterations in energy metabolism. Furthermore, study has reported that PPARα activation alleviated iron overload-induced ferroptosis in mouse liver by enhancing GPX4^[15]^. However, no study has reported the relationship between PPARα and GPX4 in the heart. PPARα is involved in cellular fatty acid uptake, esterification, and trafficking, and it also modulates genes involved in lipoprotein metabolism^[32]^. In our study, we observed a reduction in the expression level of PPARα both *in vitro* and *in vivo*, which has also been found in other studies^[33, 42]^. In the present experiment, increments of FFA and A-CoA, and reduction of ATP were observed both in vitro and in vivo after hypoxia treatment. These findings suggest reduced FFA utilization and decreased FAO under hypoxia. A-CoA produced from the oxidation of fatty acids and glucose, subsequently enters the tricarboxylic acid cycle to generate ATP^[43]^. In this study, the HH group showed increased A-CoA levels compared with the Control group, indicating reduced myocardial utilization of A-CoA, which ultimately led to lower ATP levels in the myocardium.

PPARα plays a crucial role in maintaining the balance of myocardial energy metabolism by regulating downstream genes. The induced expression of PPARα may affect the expression of CPT1 and PDK4^[44, 45]^. PDK4 is a negative regulator of glucose oxidation^[46]^. The expression of PDK4 protein decreases, and this change promotes the oxidative utilization of glucose with a corresponding decrease in the heart’s dependence on fatty acids as an energy source^[47]^. CPT1 can promote the entry of fatty acids into the mitochondrion for fatty acid oxidation^[48]^. Reduced expression of PPARα leads to decreased levels of CPT1, which in turn impairs the ability of fatty acids to enter mitochondria for oxidation^[49]^. And it was reported that PPARα can be involved in ferroptosis by mediating lipid remodeling^[50]^. SLC7A11 is a component of the cystine/glutamate antiporter (xCT), which imports cystine for glutathione biosynthesis and antioxidant defense^[51]^. SLC7A11 deficiency promotes ferroptosis by reducing glutathione synthesis^[52]^. GPX4 converts lipid hydroperoxides to lipid alcohols, thereby preventing the iron (Fe2+)-dependent formation of toxic lipid ROS^[6]^. Inhibition of GPX4 function leads to lipid peroxidation and can result in the induction of ferroptosis. In order to elucidate the mechanisms through which PPARα regulates cardiac function in this context, we applied WY14643, an agonist of PPARα, in H9c2 cells. Our results indicate that CPT1, PDK4, SLC7A11and GPX4 protein levels are elevated after PPARα activation. Our study suggests that myocardium may inhibit CPT1 and PDK4 expression by downregulating PPARα under HH, subsequently reducing mitochondrial fatty acid uptake and oxidation. Downregulation of PPARα decreased the expression of SLC7A11 and GPX4, inhibiting cystine uptake in cardiomyocytes and resulting in reduced intracellular glutathione synthesis, increased peroxide, and ROS generation, eventually leading to ferroptosis. This may be the cause of cardial injury under HH.

We next explored the causes of the PPARα changes induced by HH. Recently, it was reported that SIRT1 activation promotes the interaction between PGC1α and PPARα, which subsequently enhances mitochondrial fusion in diabetic hearts^[53]^. SIRT1 is a nicotinamide adenosine dinucleotide-dependent (NAD+) deacetylase involved in regulating energy homeostasis^[54]^. Therefore, we hypothesized that SIRT1 plays a role in regulating PPARα under hypoxia. We observed that overexpressing SIRT1 in H9c2 cells improved the hypoxia-induced changes in fatty acid oxidation and ferroptosis. Moreover, the addition of the PPARα inhibitor GW6471 inhibited the effect of SIRT1 overexpression. *In vivo* experiments showed the same effect by orally administering resveratrol, a natural agonist of SIRT1, to animals.

## 5. Conclusion

In conclusion, our study highlights the role of ferroptosis in myocardial energy metabolism changes under hypoxia, suggesting its significant association with hypoxia-induced cardiac injury. Furthermore, we explored the potential therapeutic avenue of improving hypoxia-induced alterations in cardiac fatty acid oxidation and ferroptosis through modulation of the SIRT1-PPARα-GPX4 pathway. This opens new research directions for understanding and addressing cardiac injury under hypoxic conditions.

## Declarations

## Availability of data and materials

All data generated or analyzed during this study are available from the corresponding author upon reasonable request.

